# Decipher macrophage-fibroblast-cardiomyocyte signaling interactions associated with heart failure using deep graph neural network models and single-cell RNA-seq data

**DOI:** 10.1101/2022.11.02.514928

**Authors:** Wenyu Li, Jiarui Feng, Philip Payne, Yixin Chen, Fuhai Li

## Abstract

Heart failure is a major cause of mortality. In these recent studies, novel evidence, using single-cell transcriptomic data, was reported to indicate that macrophages and fibroblasts play important roles in heart failure. The involvement of macrophages in inflammation and the relationship between inflammation and fibrosis has been established. However, the underlying molecular targets and signaling pathways mediating signaling interactions among macrophages, fibroblasts, and cardiomyocytes remain unclear. In this study, analyzing the scRNA-seq datasets using deep learning models, we ranked the cell-type specific molecular targets; and uncovered the dysfunctional intra- and inter-cellular signaling pathways that are potentially associated with heart failure. The signaling targets and pathways could be helpful in identifying effective medications for heart disease inflammation management to prevent heart failure.

## Introduction

Dilated Cardiomyopathy (DCM) is a common cardiac diagnosis that might result in heart failure. It is a major cause of morbidity despite advancement in patient care^1^. Recently, a few studies have been reported to investigate the DCM disease microenvironment (ME). For example, macrophage and signaling interactions between cardiomyocytes and macrophage are reported to be related to the cardiac diease^2,3^. The results indicated that the intricate mechanism of macrophage is dependent on its different phenotypic states. However, the underlying molecular targets and mechanisms, as well as its role in heart inflammation have not been well understood.^4^ In general, the “M1-M2 paradigm” is a ubiquitous classification of macrophage. Classically activated macrophages (M1) mediate defense of the host from a variety of bacteria, protozoa and viruses, and have roles in pro-inflammation function. Alternatively activated macrophages (M2) have anti-inflammatory function and regulate wound healing.^5^ Some studies also indicated that the polarization of macrophages can be seen as “tissue-resident vs migrated”: tissue-resident macrophages are usually regarded as

“M2-like,” however, the distinction between resident macrophages and recruited inflammatory macrophages has not been made.^6^ In general, M1 initiates inflammatory responses, mediates inflammation and activates cell defense mechanism, whereas M2 is in charge of post-inflammation damage control.

Fibroblasts are known as the non-inflammatory cell that, together with macrophage, activate and regulate the immune responses by activating pro-inflammatory signaling pathways. They are vital to pathological processes, and their phenotypes differ based on the disease type and the tissue of origin; however, our understanding of their phenotypes are often impeded by their intrinsic heterogeneity and a lack of robust subpopulation markers^7^. It has already been established that fibrosis, such as renal fibrosis, can be an end result of chronic inflammatory reactions. Similarly, chronic inflammatory reactions are strongly associated with tumourigenesis as well, insinuating a link between cancer and inflammation^8^.

Single cell RNA sequencing (scRNA-seq) has become a powerful and popular technique to measure transcriptomic variations of individual cells within disease microenvironment (ME),, which gives it great potential in discovering the regulatory mechanisms specific to a cell’s phenotype variation.^9,10^ However, it remains an open problem to rank the cell-type specific key signaling targets and infer the core signaling pathways using the scRNA-seq data. Over the years, neural networks have shown great potential in discovering heterogeneous information, novel pathogenesis, and casual genes in chronical disease progression^11–13^. Interpretable neural network models can even directly affect medical treatment.^14^ Specifically, the graph neural network (GNN), famous for its node and graph representation and classification tasks, has shown great performance on image and text data. In this study, we demonstrate how a transformer-based Graph Neural Network (GNN), named PathFinder (Feng, et al.), can discover cell type specific signaling pathways, thus help understand the dysfunctional signaling pathways in macrophage subpopulations and fibroblast cells within the disease ME of Dilated Cardiomyopathy (DCM).

## Results

### A: Cell type classification accuracy

In the interest of the robustness of our methods results, only 8192 of the highest expressed genes are fed into the network. The pre-defined pathways are generated from the combined database consisting of ligand-receptor network, signaling network, and gene-regulation network. The Pathfinder model is trained on the receptor cell expression matrix first, and the intra-cell network is generated as a result of ranking all learnable weights of the resulting pathways. Note that in this study we selected up-regulated pathways as output. **Fig 1** shows a general flow of our method. The model is applied on macrophage subpopulation 1 (M1), macrophage subpopulation 2 (M2), fibroblasts and cardiomyocytes in 45 patients’ samples. The model takes 2 datasets as input, control vs test. The following experiments are then conducted such that healthy datasets are the control group and diseased (DCM) cells are the test group. In other words, the direct inputs to the network are 8192 genes and the expression matrices of size *cell count for healthy* x 8192 and *cell count for DCM* x 8192. (For details of the experiments, see Method section) As shown in the **Table 1**, PathFinder correctly predicts most of the conditions of each cell, and generates intra cell communication networks for M1, M2, fibroblast and cardiomyocytes

**Table 1:** Performance of Pathfinder on all cell states (networks used for plotting inter-cell networks are highlighted as **bold**).

|  | **ACC** | **Recall** | **Precision** | **Specificity** | **F1** | **AUC** |
| --- | --- | --- | --- | --- | --- | --- |
| M1 | 0.72 | 0.22 | 0.80 | 0.97 | 0.35 | 0.68 |
| **M2** | 0.83 | 0.89 | 0.89 | 0.59 | 0.89 | 0.83 |
| **M1 with resampling** | 0.73 | 0.44 | 0.91 | 0.97 | 0.6 | 0.8 |
| M2 with resampling | 0.66 | 0.71 | 0.65 | 0.62 | 0.68 | 0.74 |
| **Fibroblast** | 0.71 | 0.63 | 0.75 | 0.79 | 0.69 | 0.78 |
| Fibroblast with resampling | 0.69 | 0.55 | 0.77 | 0.83 | 0.64 | 0.77 |
| Cardiomyocyte | 0.88 | 0.43 | 0.57 | 0.95 | 0.49 | 0.82 |
| **Cardiomyocyte with resampling** | 0.89 | 0.79 | 0.97 | 0.98 | 0.87 | 0.92 |

### B: The up-regulated signaling targets are identified in the top-ranked signaling pathways

To validate our model’s integrity on learning the most important difference of the two conditions, like normal control vs DCM, we extract the top 200 pathways selected by Pathfinder and validate our model against the Differentially Expressed Genes (DEG) analysis. Each upregulated pathway consists of multiple genes, whose average log2 fold change over gene features are calculated by FindMarkers from Seurat^15^, then we take the average of all genes’ fold change values as the final fold change for the corresponding pathway. Note that positive fold change means that the gene is up-regulated in test group. The notation for the final log2 fold change *C^p^* is shown below:

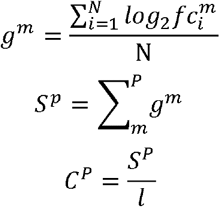

where N is the total number of cells in each macrophage subpopulation, w represents a cell type and m represents a specific gene and l is the number of genes in pathway P. *S^p^* is the sum of all log2 fold change values for the genes in a specific pathway P. The mean of total log2 fold change, 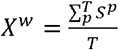, where T is the total number of pathways identified by Pathfinder, is 0.06 for Mac1 and 1.23 for Mac 2, which suggests that Mac1 paths are only slightly up-regulated while Mac2 paths are mostly significantly up-regulated in diseased cells. The distributions of the top 200 pathways selected by Pathfinder and the rest are shown in **Figs 2a-d**, where the average gene log2 fold change value *C^p^* is used. For ease of comparison, we take the density of the distribution.

**Fig 2.**
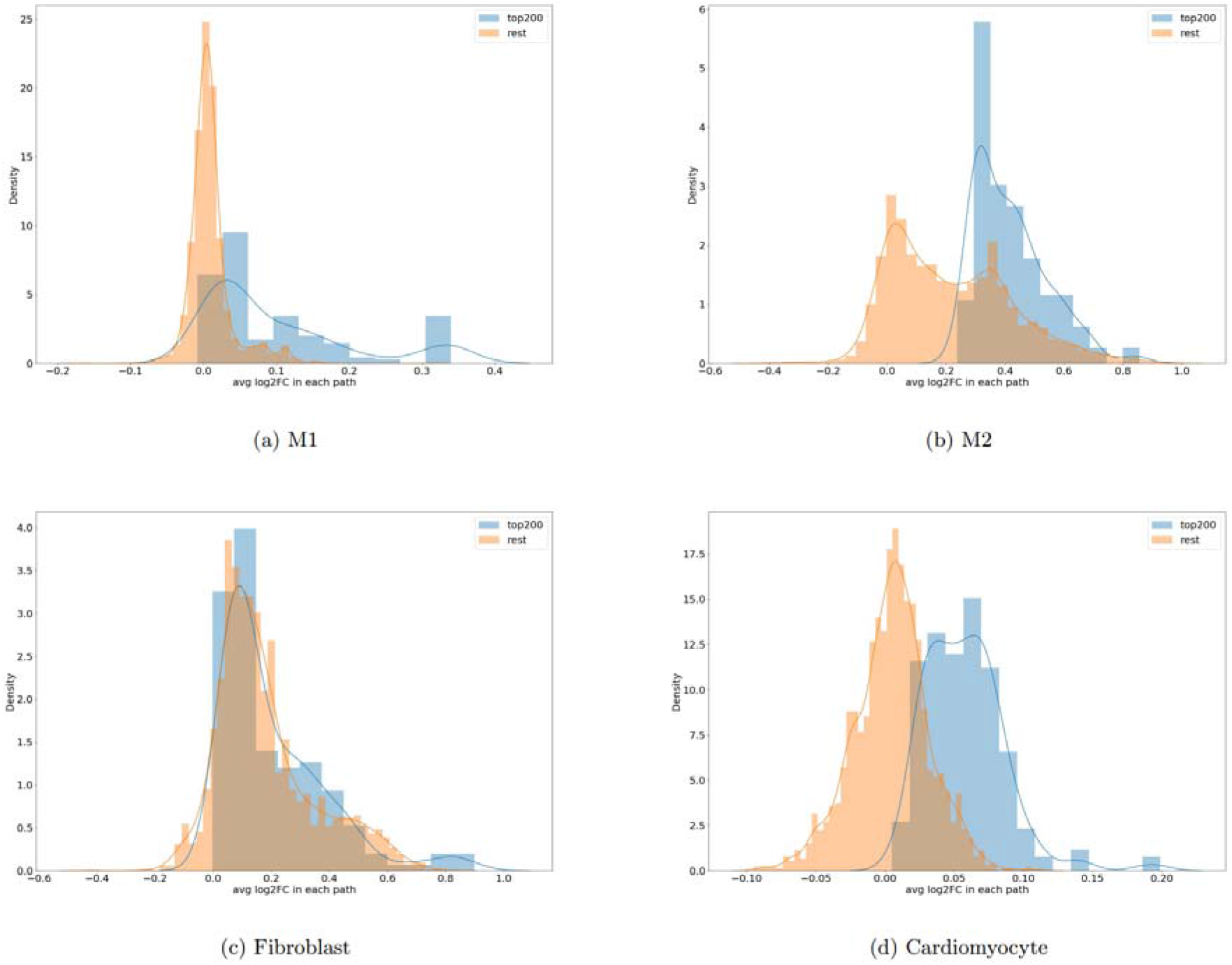
Top 200 paths and the rest total log2 fold change distribution comparison.

As seen that Pathfinder is able to identify up-regulated signaling pathways for all cell types. The distribution of the top 200 paths is skewed to the right and almost all the paths have a positive total log2 fold change. Even though there’re negative pathways selected by the model, it could mean they are also important for distinguishing the two groups. The results validated Pathfinder’s efficiency and the correctness.

### C: Dysfunctional intra-cellular signaling pathways of macrophages, fibroblasts and cardiomyoctes

To uncover the underlying mechanism of inflammation in DCM, the intra-cellular networks are generated from the pre-defined path lists, which are defined as the shortest path between each two genes and all possible paths from the receptor to the target gene. The network is of the top 200 highly represented paths in the training process. Note that the ligand-receptor interactions are excluded from the intra-networks. The intra networks for M1, M2, fibroblast and cardiomyocyte are shown in **Figs 3–6**.

**Fig 3.**
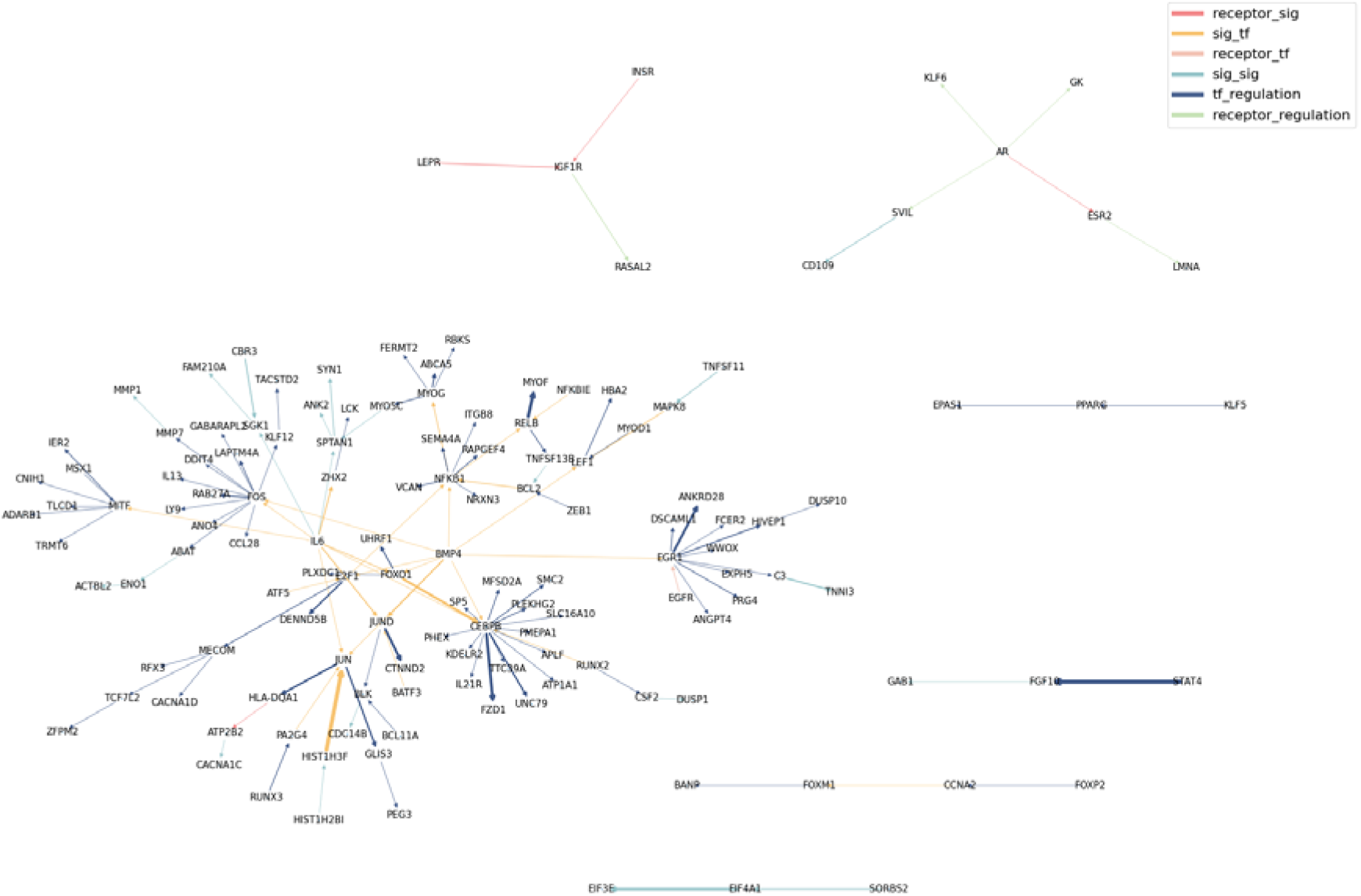
Dysfunctional intra-cellular signaling networks of M1. Edge width indicating pathway weights.

The intra-cellular signaling network for Mac1 consists of 4 receptor-signaling interaction pathways, 29 signal-transcription factor interaction pathways, 1 receptor-transcription factor interaction pathways, 21 signal-signal interactions, 78 tf-regulation interactions and 5 receptor-regulation interactions.

**Fig 3.**
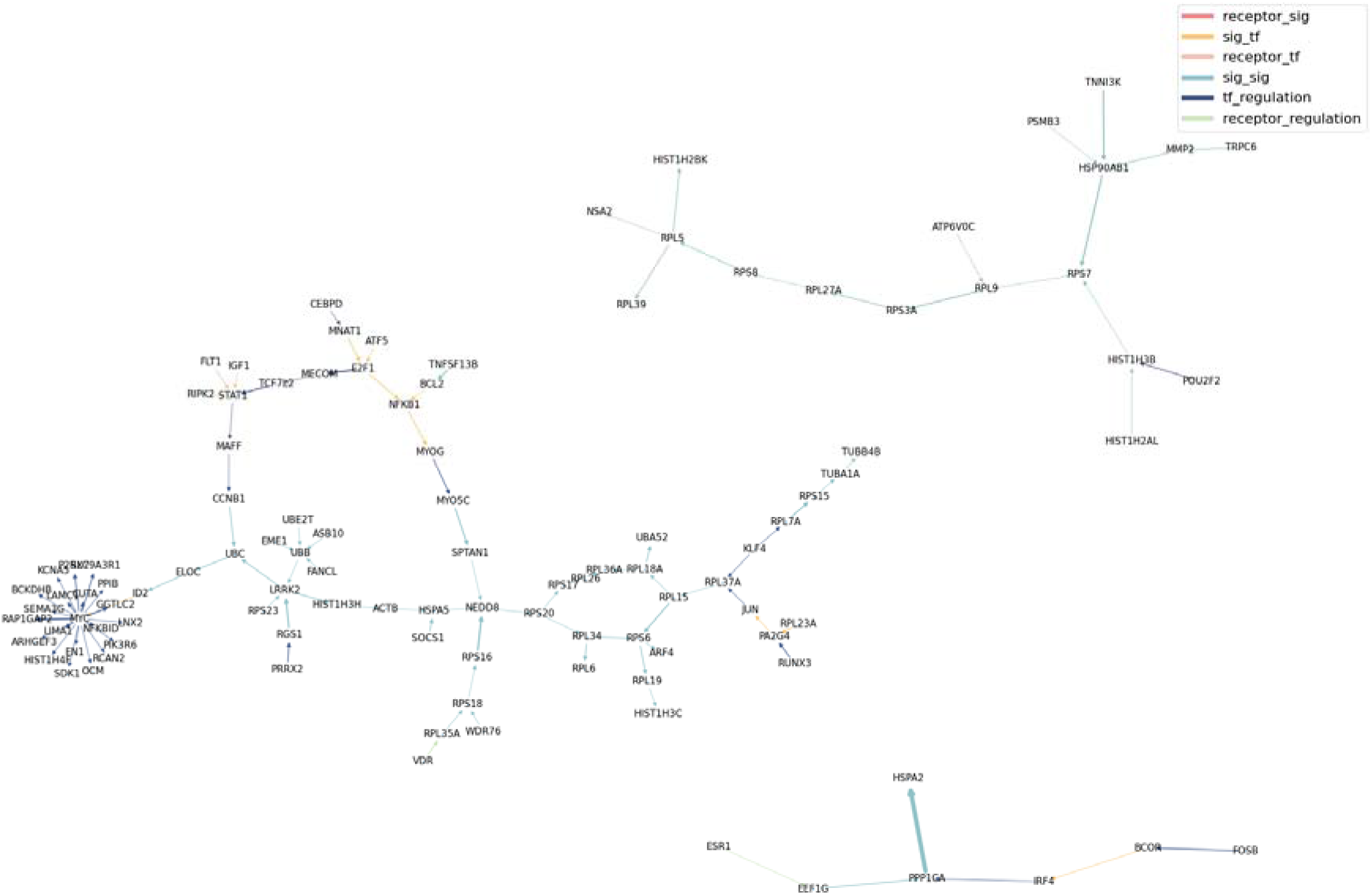
Dysfunctional intra-cellular signaling networks of M1. Edge width indicating pathway weights.

The final intra network for Mac2 consists of 5 receptor-signaling interaction pathways, 27 signal-transcription factor interaction pathways, 3 receptor-transcription factor interaction pathways, 66 signal-signal interactions, 66 tf-regulation interactions and 16 receptorregulation interactions.

**Fig 3.**
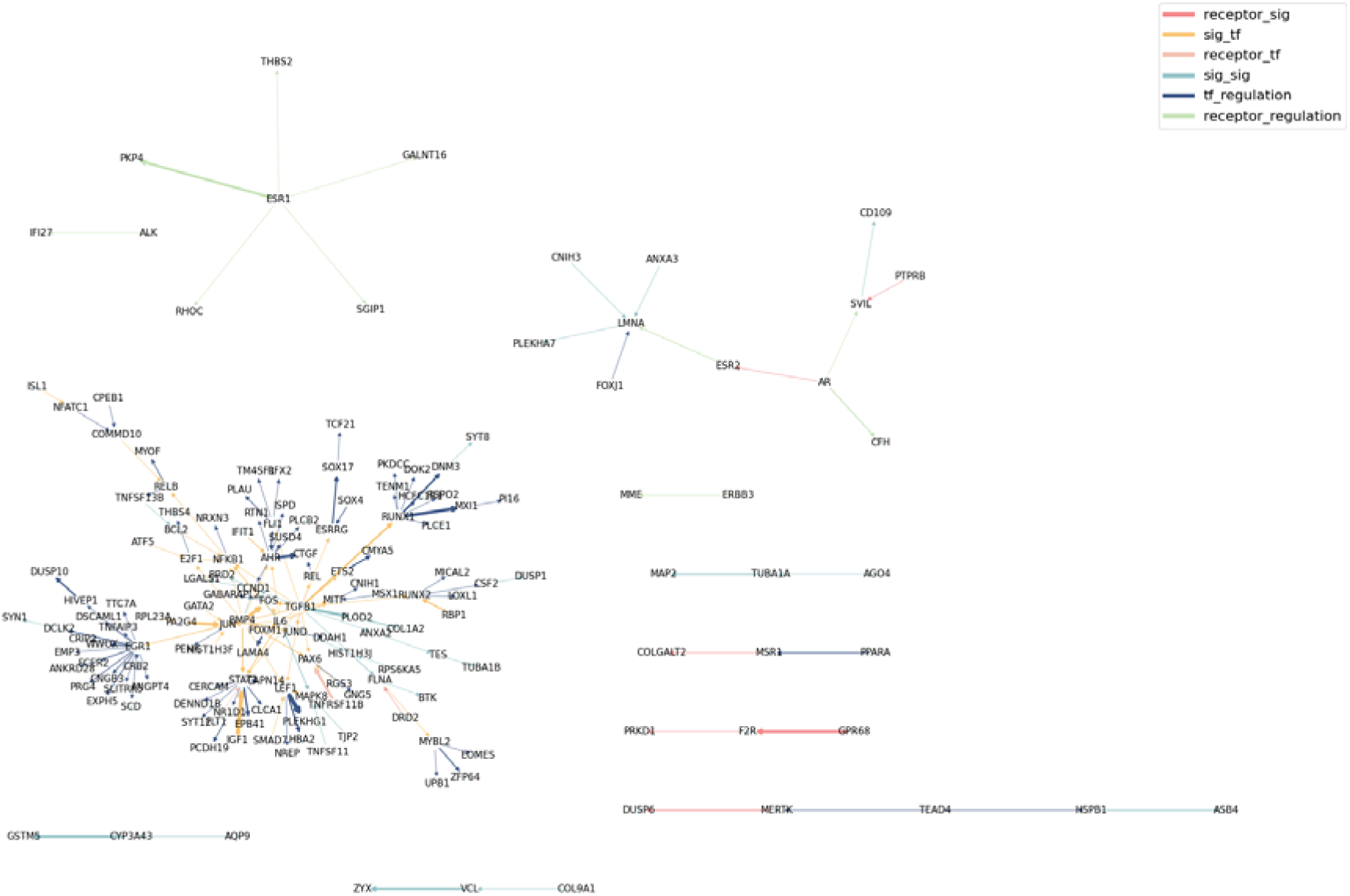
Dysfunctional intra-cellular signaling networks of Fibroblasts. Edge width indicating pathway weights.

The final intra network for fibroblast consists of 7 receptor-signaling interaction pathways, 51 signal-transcription factor interaction pathways, 2 receptor-transcription factor interaction pathways, 33 signal-signal interactions, 76 tf-regulation interactions and 10 receptor-regulation interactions.

**Fig 3.**
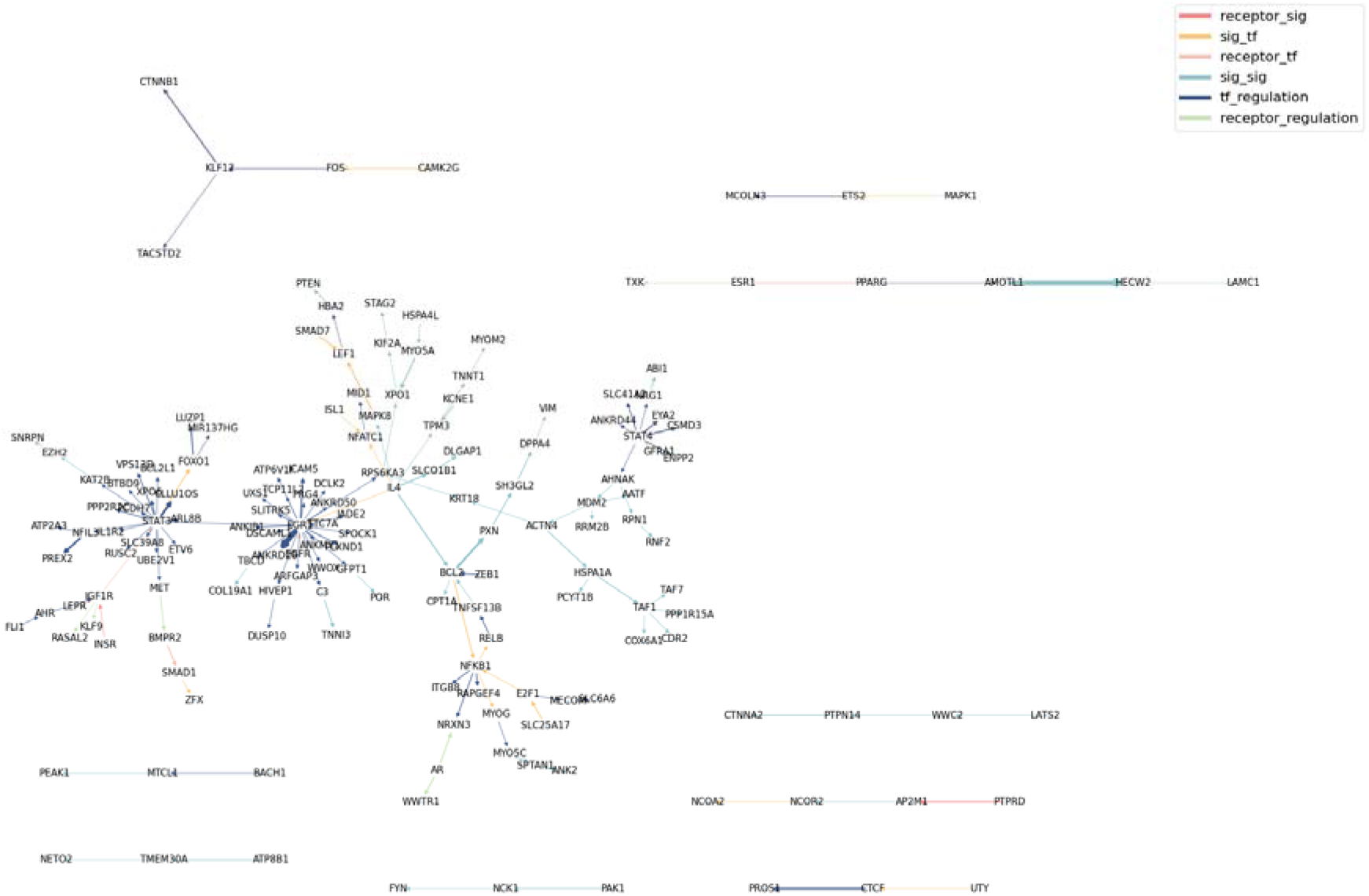
Dysfunctional intra-cellular signaling networks of cardiomyocyte. Edge width indicating pathway weights.

The final intra network for fibroblast consists of 3 receptor-signaling interaction pathways, 16 signal-transcription factor interaction pathways, 5 receptor-transcription factor interaction pathways, 55 signal-signal interactions, 71 tf-regulation interactions and 6 receptor-regulation interactions.

### D: Macrophage-Fibroblast-Cardiomyocte inter-cellular signaling interactions

To understand the interactions between cardiomyocytes, macrophage and fibroblasts, the inter-networks are generated as follows: The intra cell communication networks for ligand and receptor cells need to be generated first, then, the ligands of ligand cell and receptors of receptor cell will be extracted from the intra network respectively. Finally, ligand-receptor pairs are based on the ligand-receptor database. The difference between inter-cell networks and intra-cell networks are the addition of ligand-receptor interactions. The inter-networks between M1, M2 and fibroblast are shown in **Fig. 7–9.** From the results, *BMP4* is present in M1 and fibroblast inter cellular interactions. It is found to induce inflammation in certain stages of disease.^16^ *IGF1* has interactions with cardiomyocyte in both fibroblast and M2. It is shown in study that IGF1 is related to muscle regeneration^17^

**Fig 7.**
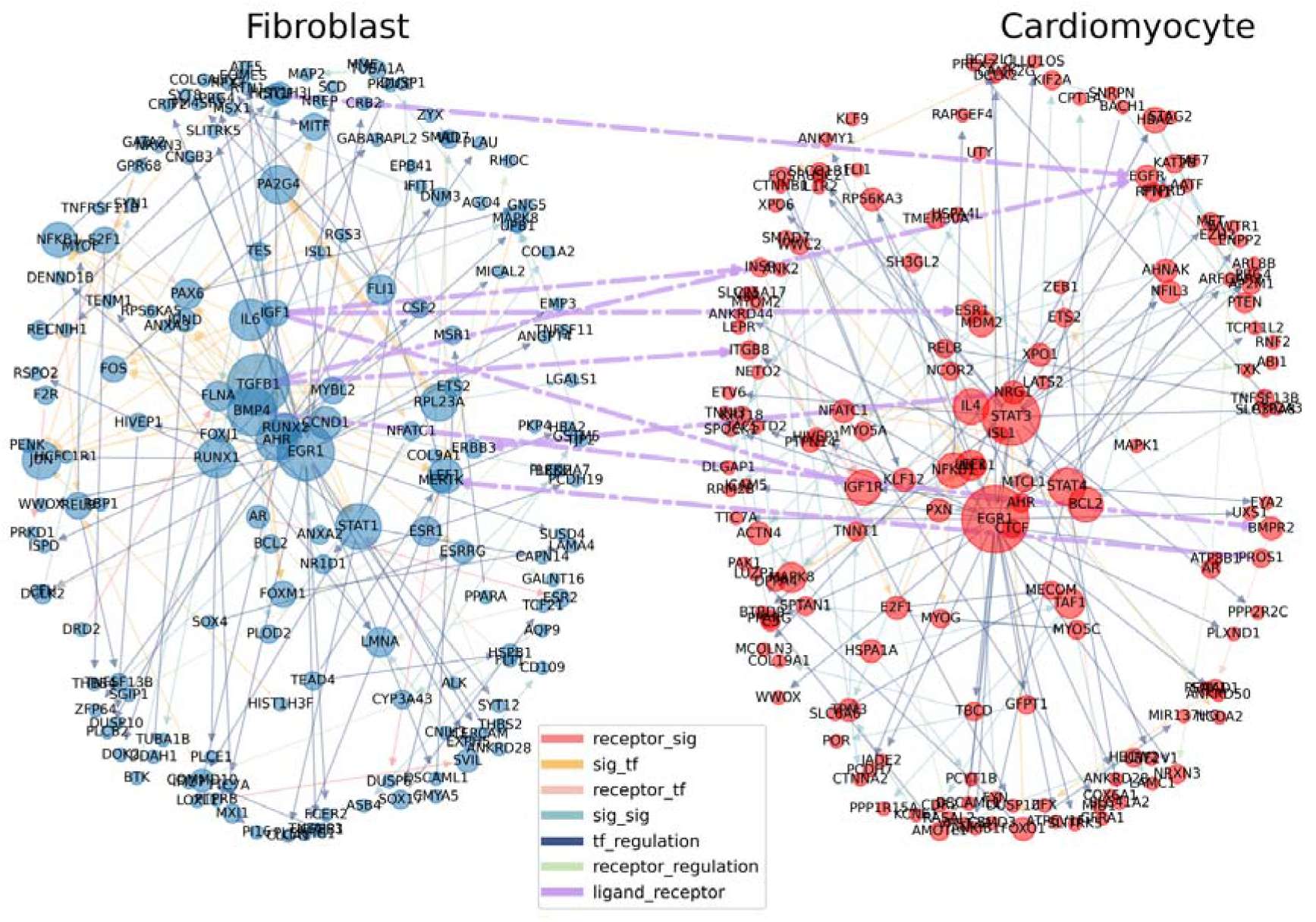
Inter network of fibroblast and cardiomyocyte, with the node size and edge width being the weight of the edge for the corresponding cell type.

**Fig 8.**
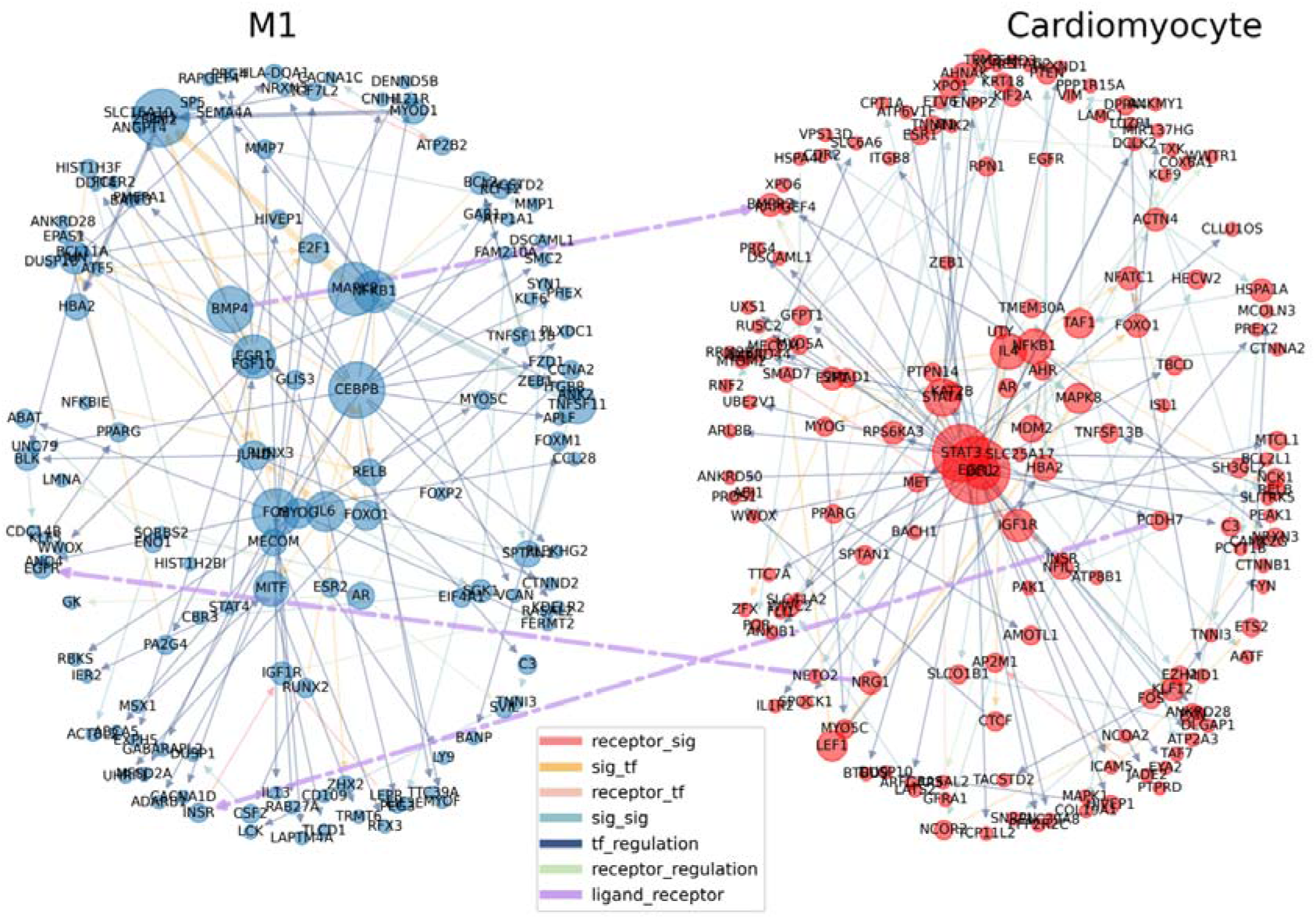
Inter network of Mac1 and cardiomyocyte, with the node size being the weight of the edge for the corresponding cell type.

**Fig 9.**
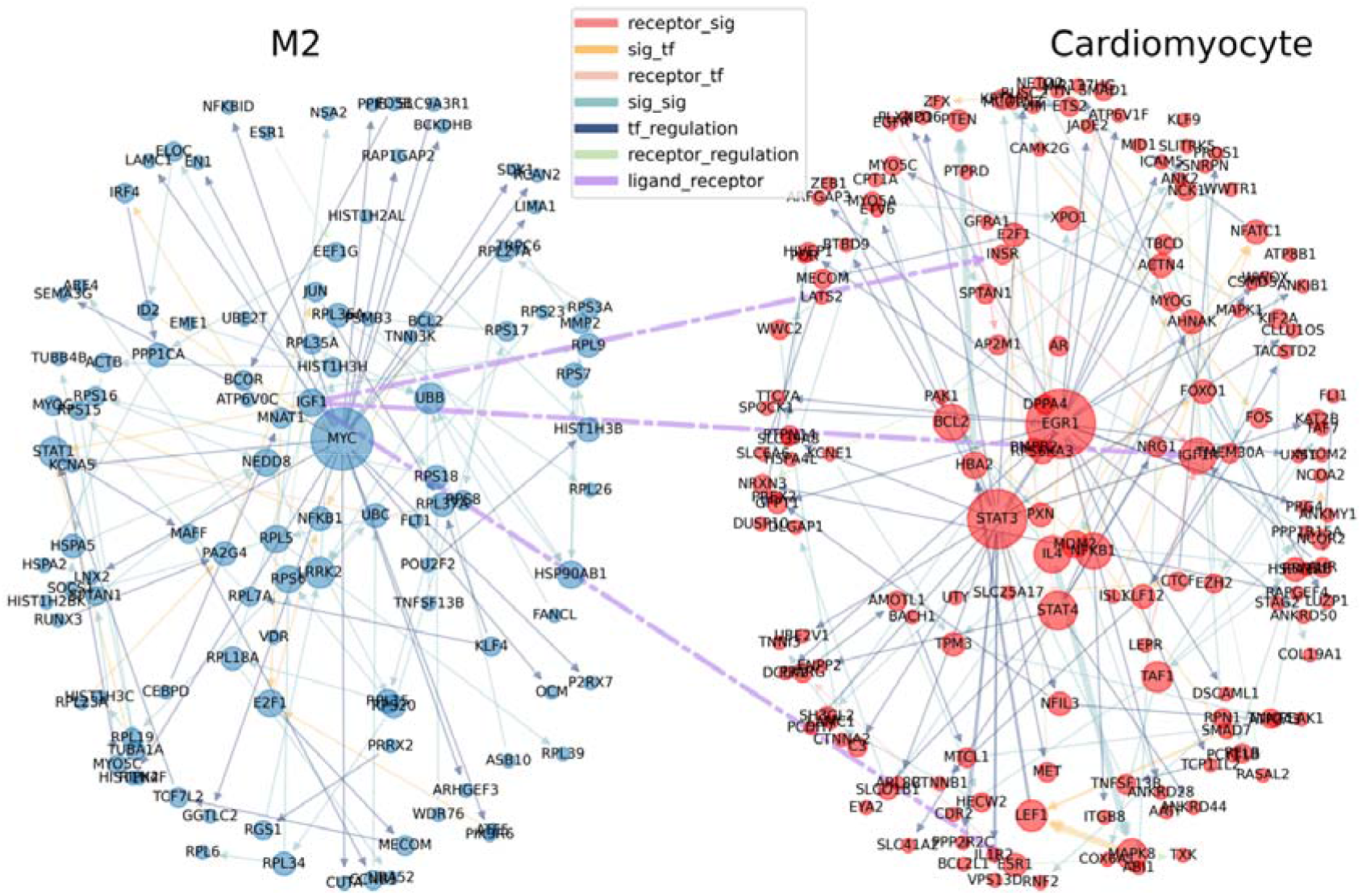
Inter network of Mac2 and cardiomyocyte, with the node size being the weight of the edge for the corresponding cell type.

### KEGG pathway enrichment analysis

To further test the model’s ability to find important signaling pathways, we extracted KEGG pathways from the previous results. The KEGG pathways on individual states of macrophage reveal multiple immune cell differentiations, and M1 and fibroblast revealed an enrichment in inflammatory pathways, which confirms the model’s correctness in analyzing gene-gene interactions.

KEGG analysis is conducted on the genes in the top 500 pathways found by Pathfinder and those found by the previously mentioned statistical selection. The resulting number of pathways is shown in **Table 5**.

**Table 5.** Number of KEGG pathways identified.

|  | M1 | M2 | Fibroblast |
| --- | --- | --- | --- |
| Pathfinder | 244 | 208 | 238 |
| Statistical | 122 | 289 | 272 |

After filtering all pathways by adjusted p value < 0.05, the number of pathways left in both methods are shown in **Table 6**:

**Table 6.** Number of KEGG pathways identified after filtering.

|  | M1 | M2 | Fibroblast |
| --- | --- | --- | --- |
| Pathfinder | 122 | 46 | 101 |
| Statistical | 6 | 74 | 22 |

Note that the number of pathways for statistical selection shrinks significantly, especially for M1 and fibroblasts, which means that pathways Pathfinder extracted are more enriched in general. The KEGG analysis results in an adjusted p value and log 2 FC scores for each pathway found in the database. We choose adjusted p value as an indicator for enrichment score. To make the number more intuitive, we take the negative log2 of the adjusted p value to make the results positively proportional to the enrichment level. The overlapping and unique pathways in each cell type is plotted in **Fig 10**. We can see that Pathfinder finds much more pathways than Seurat for M1, less pathway for M2 and more pathways for fibroblast. Then the pathway enrichment results for all Pathfinder experiments are obtained by merging all cell types in **Fig 11**. We use 0 to indicate absence in the corresponding cell type and rescale the scores to range 0 to 1.

**Fig 10.**
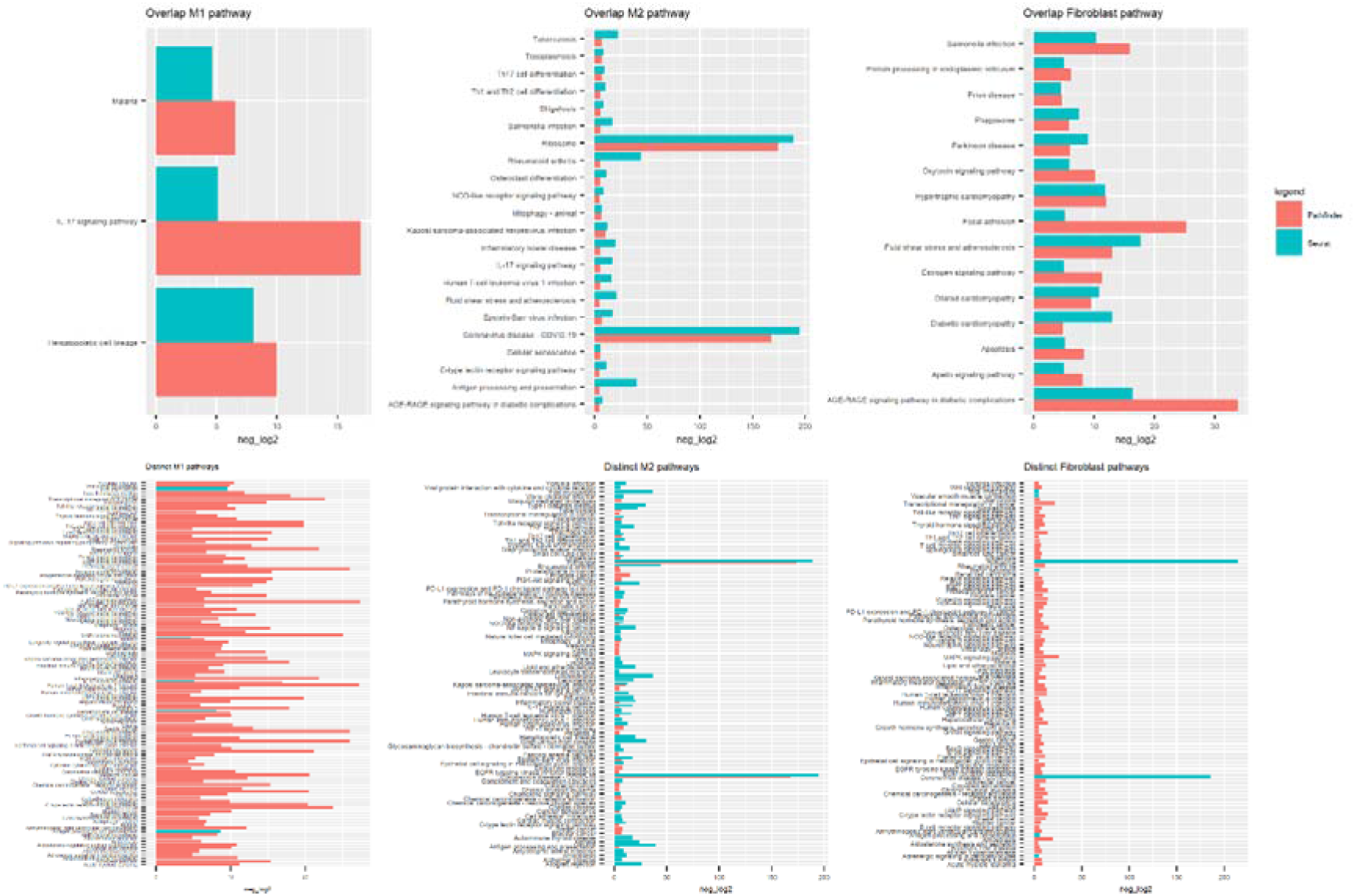
-log2 of the adjusted p value of the pathways found by both Pathfinder and statistical selection (annotated by “Seurat”)

**Fig 11.**
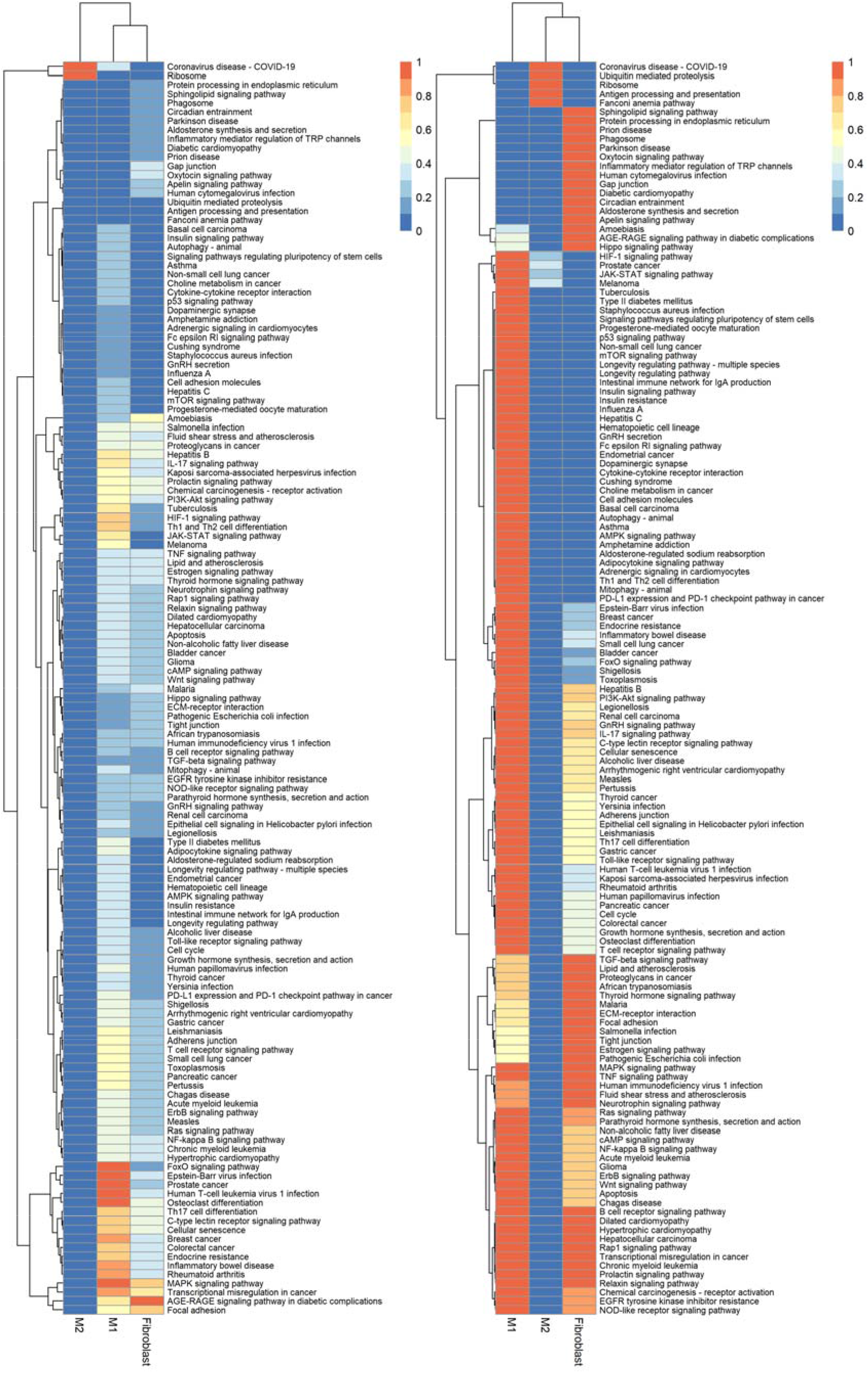
KEGG pathway for M1, M2 and Fibroblast, with scores rescaled across rows (left) and across columns (right)

While general Differentially Expressed Genes analysis extracted many genes, Pathfinder is able to find several key genes that are typical to inflammatory processes. Normalization across cell types reveals an enrichment in inflammatory pathways in both M1 and fibroblast, and since the network is designed to learn up-regulated paths in diseased samples, the fact that DCM pathway is most enriched in M1 indicates that M1 may be pro-inflammatory compared to M2. Apart from the obvious heart-related disease pathways like dilated cardiomyopathy, many other pathways, such as NF-kappa B activation, are more enriched in M1, which corresponds to the belief that one of M1’s most important roles is initiation and resolution of inflammation.^18^ NF-kappa B is also enriched in fibroblast. It is a transcriptional factor widely known for its potency to activate transcription of many genes encoding immunologically relevant proteins.^19^ NF-kappa B also generates a loop that maintains the inflammatory signals as it controls the expression of genes encoding pro-inflammatory factors, which are all very important for controlling the inflammation process. Due to the strong connection between inflammation and fibrosis, inhibiting the activation of NF-kappa B also controls the progression of chronic kidney disease.^8^ Other indicators of pro-inflammatory characteristics exist in KEGG pathways in M1 and fibroblast, like MAPK signaling, Th1 and Th2 cell differentiation (stimulate strong cell-mediated immune responses), IL-17 signaling (an inflammation-mediated pathway). Those pathways are not emphasized in the statistical analysis results, which proves that the DL model is able to recognize key genes/paths that can’t be picked out by DEG analysis. Cell damage generally activates MAPK signaling pathway, which directs inflammatory response to a series of stimuli, including inflammatory cytokines. Study has also shown that certain genes can down-regulate M1 polarization by inhibiting the MAPK pathway.^20^ M1 can cause tissue damage by facilitating Th1 response, and M2 can maintain tissue integrity by promoting the Th2 response.^21^ Th17 Cell differentiation also plays an important role in the pathogenesis of inflammatory and autoimmune diseases.^22^

Multiple inflammatory diseases, including Inflammatory bowel disease, Human T-cell leukemia virus 1 infection, Epstein-Barr virus disease, other infections and cancer are comparatively more enriched in M1 and fibroblast.

## Method

### snRNA-seq and scRNA-seq dataset

In this work, the snRNA-seq and scRNA-seq datasets are collected from Gene Expression Omnibus (GEO) database with accession number GSE183852^1^. Their heart specimens are obtained from healthy donors and patients with Dilated (nonischemic) Cardiomyopathy. 38 samples (25 healthy, 13 diseased) are collected as snRNA-seq, while 7 samples (2 healthy, 5 diseased) are collected as scRNA-seq using the 10X Genomics 5□ Single Cell platform. Both single-nucleus and single-cell libraries were sequenced and aligned, then filtered for quality control (QC) including unsupervised clustering, PCA and UMAP^23^ reduction and differential expression analysis using R packages Harmony and Seurat^15^. The final integrated dataset consists of 220,752 nuclei and 49,723 cells representative of 14 major cell types. Details are shown in **Table 2**.

**Table 2:** Distribution of major cell types in integrated dataset.

|  | **DCM** | **Donor** | **Perc %** | **Total** |
| --- | --- | --- | --- | --- |
| **Cardiomyocytes** | 6478 | 40620 | 17.457 | 47098 |
| **Fibroblasts** | 36574 | 35593 | 26.749 | 72167 |
| **NK/T-Cells** | 4597 | 2768 | 2.73 | 7365 |
| **Macrophages** | 14750 | 19994 | 12.878 | 34744 |
| **Smooth Muscle** | 1600 | 3399 | 1.853 | 4999 |
| **Lymphatic** | 1027 | 1285 | 0.857 | 2312 |
| **Neurons** | 923 | 1239 | 0.801 | 2162 |
| **Endothelium** | 20923 | 28471 | 18.308 | 49394 |
| **Epicardium** | 466 | 933 | 0.519 | 1399 |
| **Mast** | 206 | 1125 | 0.493 | 1331 |
| **Adipocytes** | 365 | 706 | 0.397 | 1071 |
| **B-Cells** | 253 | 92 | 0.128 | 345 |
| **Pericytes** | 9038 | 20267 | 10.862 | 29305 |
| **Endocardium** | 5957 | 10145 | 5.968 | 16102 |
| **Perc %** | 38.235 | 61.765 |  |  |
| **Total** | 103157 | 166637 |  | 269794 |

The dimensionality reduction plot for all cell types using UMAP is plotted in **Fig 14** with a side-by-side healthy vs DCM comparison. The clusters are drawn using resolution 0.6.

**Fig 14.**
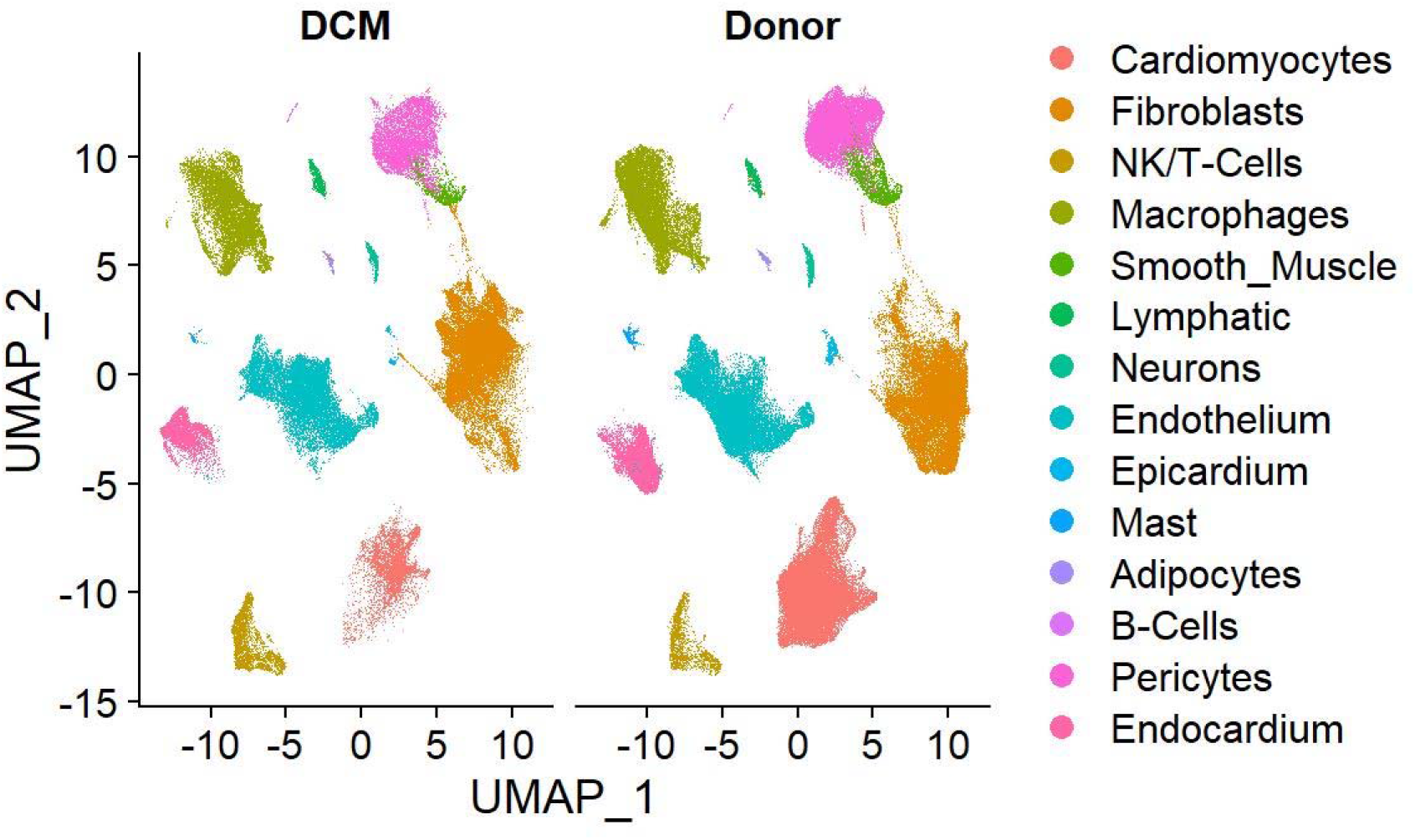
Dimension plot of all types of cells in the original sample Then the dataset is divided into subsets according to cell types.

### Macrophage

Although there’re various sets of markers to characterize macrophage phenotypes, in this work, the Mac1 subpopulation is identified by the tissue-resident markers: MRC1, SIGLEC1, CD163, LYVE1, F13A1, and Mac2 by chemokines and cytokines markers: CCL3, CCL4, CXCL3, CXCL8, IL1ß.^1^ The classification of Mac1, Mac2 and others are based on the gene expression level on RNA assay. We first conducted clustering with resolution 0.1 to divide a total of 34744 macrophages into 5 categories^1^, where M1 and M2 are identified by their markers, then we separated the diseased (DCM) samples and the healthy samples, constructing the final datasets used for the model. The corresponding markers are in the supplemental tables. The UMAP distribution of the subpopulations of macrophage is plotted in **Fig 15**.

**Fig 15.**
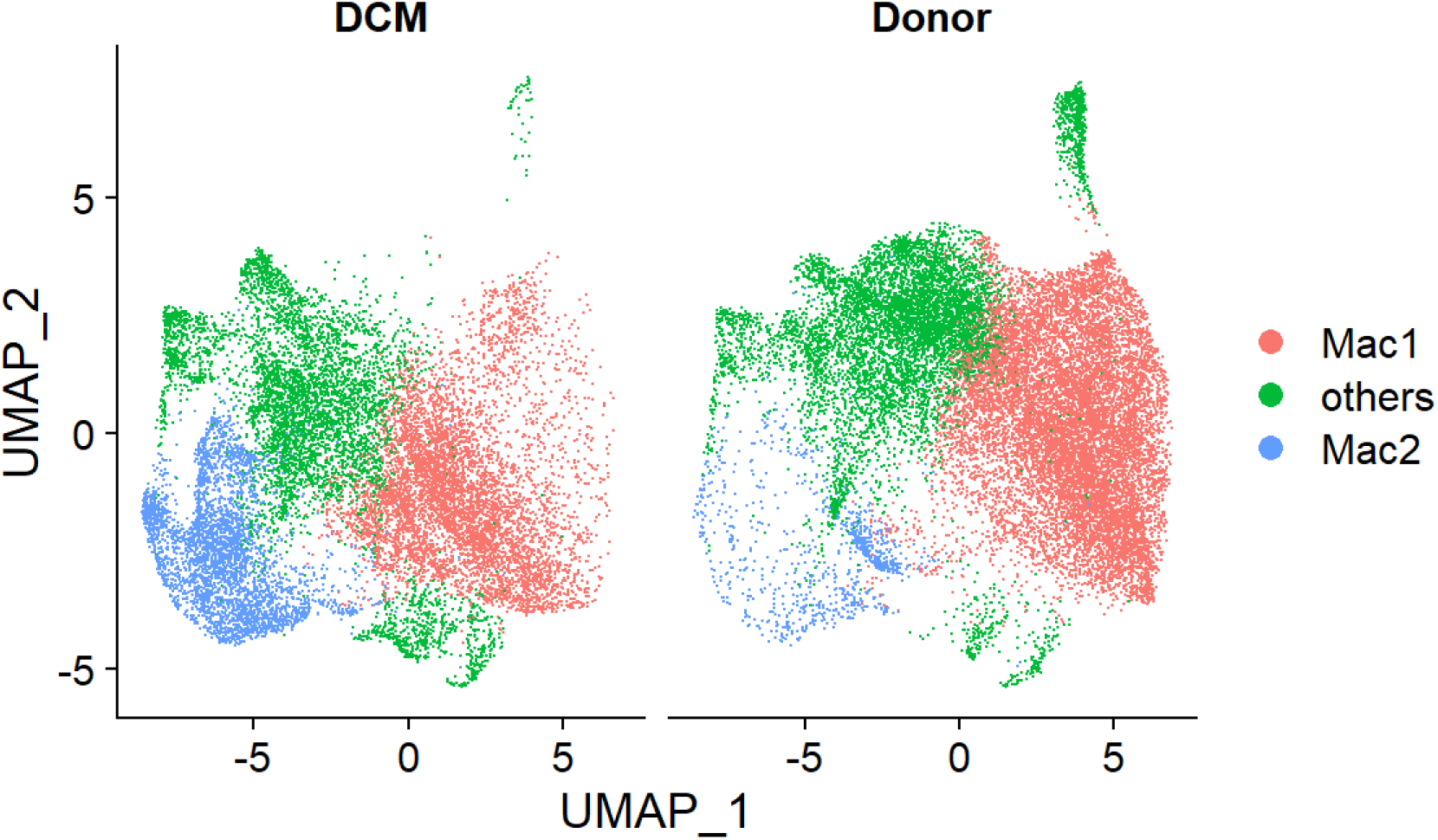
Distribution of Macrophage subpopulation with resolution 0.1, dim = 1:80.

We focus specifically on the Mac1 and Mac2 subpopulation since there’s a notable increase in Mac2 and slight decrease in Mac1 from healthy to DCM samples, which, according to the previous studies, indicate the M1 are pro-inflammatory while the M2 are anti-inflammatory^1,5^; It is further inferred that M1 is the tissue-resident macrophage subpopulation, which is confirmed by the tissue-resident markers; and M2 is the migrated subpopulation. The distribution of the data is shown in **Table 3**. Note that healthy Mac1:diseased Mac1 is around 66.4:33.6, while healthy Mac2:diseased Mac2 is around 79:21.

**Table 3:** Distribution of M1, M2 and fibroblast.

|  | M1 | M2 | Percentage | Total |
| --- | --- | --- | --- | --- |
| **DCM** | 5981 | 3520 | 42.6% | 9501 |
| **Healthy** | 11843 | 939 | 57.4% | 12782 |
| **Percentage** | 80% | 20% |  | 100% |
| **Total** | 17824 | 4459 | 100% | 22283 |

### Fibroblast

The classification of fibroblast subpopulation is based on previous study^1^: we used resolution 0.2 to divide fibroblast into 9 subtypes, representing Fb subtypes 1-8 and Epicardium. We decided to train the model on the entire population to distinguish different disease states for the two following reasons: as shown in **Fig 16–17**, the clustering results suggest that except for cluster 0 and 6, other subpopulations have obvious differences in expression in control and test groups. In addition, the lack of robustness in markers for fibroblast subpopulation might add bias to the analysis.

**Fig 16.**
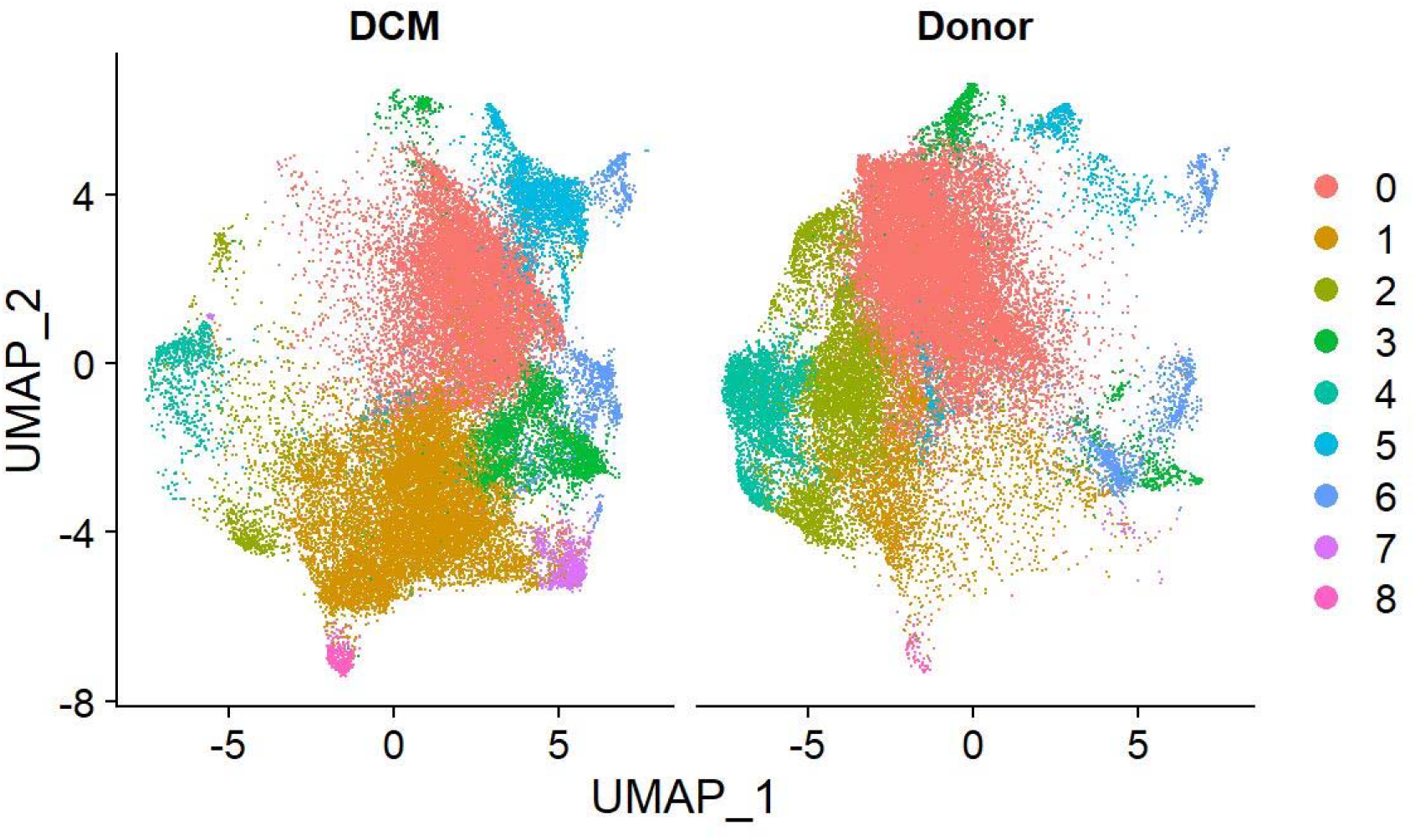
Distribution of Fibroblasts subpopulation (Fb1-8 & Epicardium) with resolution 0.2, dim = 1:80.

**Fig 17.**
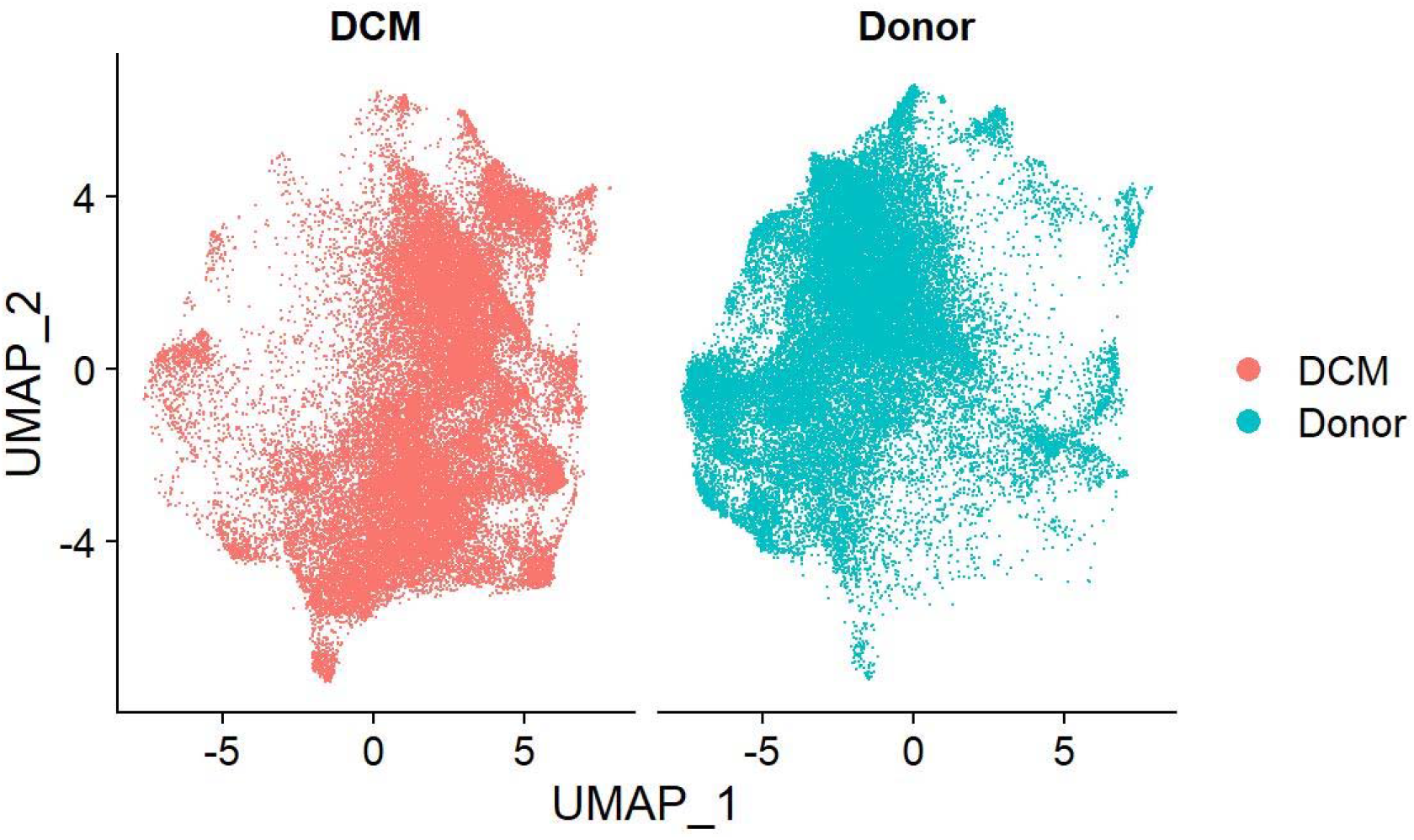
Fibroblasts dimension plot based on disease state.

A general comparison in **Fig 17** between control and test groups shows that fibroblasts have an overall difference in gene enrichment. The distribution of the input data for fibroblast is shown in **Table 5**. In the experiment, no resampling is applied due to the already very balanced data.

**Table 4.** Distribution of Fibroblast.

|  | Count | Percentage % |
| --- | --- | --- |
| DCM | 36574 | 50.68 |
| Healthy | 35593 | 49.32 |
| Total | 72167 | 100 |

To account for the problem of imbalance, we utilized the Synthetic Minority Over-sampling Technique (SMOTE)^24^, which is essentially oversampling with KNN. Specifically, it synthesizes new data in the same neighborhood for the minority class, which makes the resulting model better at generalization than randomly duplicating minority data. We used minority:majority = 8:10 for M1, and minority:majority = 1:1 for the other cell types.

### Model Structure & Notations

The PathFinder model was developed in reference (Feng, et al.). In brief, it improves the graph neural network model by using Graphormer^26^, which is designed for signaling network analysis using single cell RNAseq data.

## Conclusion

Contrary to some beliefs, the M1 phenotype characterized by resident markers show signs of pro-inflammatory interactions, so it may be the evidence that tissue-resident macrophages are not strictly anti-inflammatory. However, M2 phenotype characterized by CCL3, CCL4, CXCL3, CXCL8, IL1β are not strictly anti-inflammatory either. In conclusion, M1 is relatively more pro-inflammatory than M2, and Fibroblast has a combined effect on inflammation. There’re, as expected, also many overlapping pathways between macrophage and fibroblast, and some of them suggest that pathways induced or affected by inflammation also have a big impact on fibrosis.

## Limitations

While our model can directly generate gene-gene interaction networks with meaningful and relatively accurate results, it still suffers from the common limitations present in GNNs used for inferring gene networks: biological information embedding and effective feature engineering. So far, the Pathfinder model can’t learn the regularization term of fold change. However, in practice it may not be reasonable to set a fixed regularization term of fold change, since it should be a part of the classification learning process. Also, Pathfinder can’t study the chronical development, or multiple stages, of a disease, but in reality, it’s usually very important, and there’re probably more cell types involved in the process.

